# Quantitative modeling of regular retinal microglia distribution

**DOI:** 10.1101/2021.06.30.450549

**Authors:** Yoshie Endo, Daisuke Asanuma, Shigeyuki Namiki, Kenzo Hirose, Akiyoshi Uemura, Yoshiaki Kubota, Takashi Miura

## Abstract

1

Microglia are resident immune cells in the central nervous system (CNS), showing a regular distribution. Advancing microscopy and image processing techniques have contributed to elucidating microglia’s morphology, dynamics, and distribution. However, the mechanism underlying the regular distribution of microglia remains to be elucidated.

First, we quantitatively confirmed the regularity of the distribution pattern of microglial soma. Second, we formulated a mathematical model that includes factors that may influence regular distribution. Next, we experimentally quantified the model parameters (cell movement, process formation, and ATP dynamics). The resulting model simulation from the measured parameters showed that direct cell-cell contact is most important in generating regular cell spacing. Finally, we tried to specify the molecular pathway responsible for the repulsion between neighboring microglia.

**Author summary:** Microglia are resident immune cells in the central nervous system. It is known that the microglia cells show a regular distribution, but the mechanism underlying the regular distribution remains to be elucidated. In the present study, we quantitatively assayed the regularity of the microglia soma distribution using the image processing technique. Next, we formulated a mathematical model of cell distribution that includes factors that may influence regular distribution. Next, we experimentally quantified the model parameters using organ culture. As a result, we obtained parameters for cell migration, cell process dynamics, and extracellular ATP dynamics. Then we undertook numerical simulation of the mathematical model using the parameters obtained by the experiments. The resulting model simulations showed that direct cell-cell contact is the most important factor in generating regular spacing. Finally, we screened possible molecular pathways involved in the regular spacing of microglia, which confirmed the validity of the model.

## 2 Introduction

Microglia are resident macrophages in the CNS and show a regular distribution ([1], Fig. 1A). Microglia cells have several physiological functions; control of neuronal cell production, neural migration, axonal growth, synaptogenesis [2, 3] and angiogenesis [4]. Microglia cells also plays a role under pathological conditions; defense against infection, inflammation, trauma, ischemia, tumor, and neurodegeneration [5]. Microglia have two modes of action (Fig. 1B) - in the healthy brain, they are inactive (resting) and become active during an immune response [6]. The resting and activated microglia have distinct morphology (Fig. 1C). Resting microglia have a small soma and elongated ramified processes extending and retracting continuously, surveying their microenvironment [7]. When microglia recognize a pathogen or other inflammatory stimulus, they rapidly become an active state, retract their processes and become efficient mobile effector cells. ATP mediates the chemotaxis toward the injured site. Injured cells release ATP, activating the microglial P2X and P2Y receptors. Extracellular ATP regulates microglia branch dynamics and mediates microglial movement toward the injury [8–10]. Local injection of ATP can mimic this immediate chemotactic response [11].

**Fig 1.**
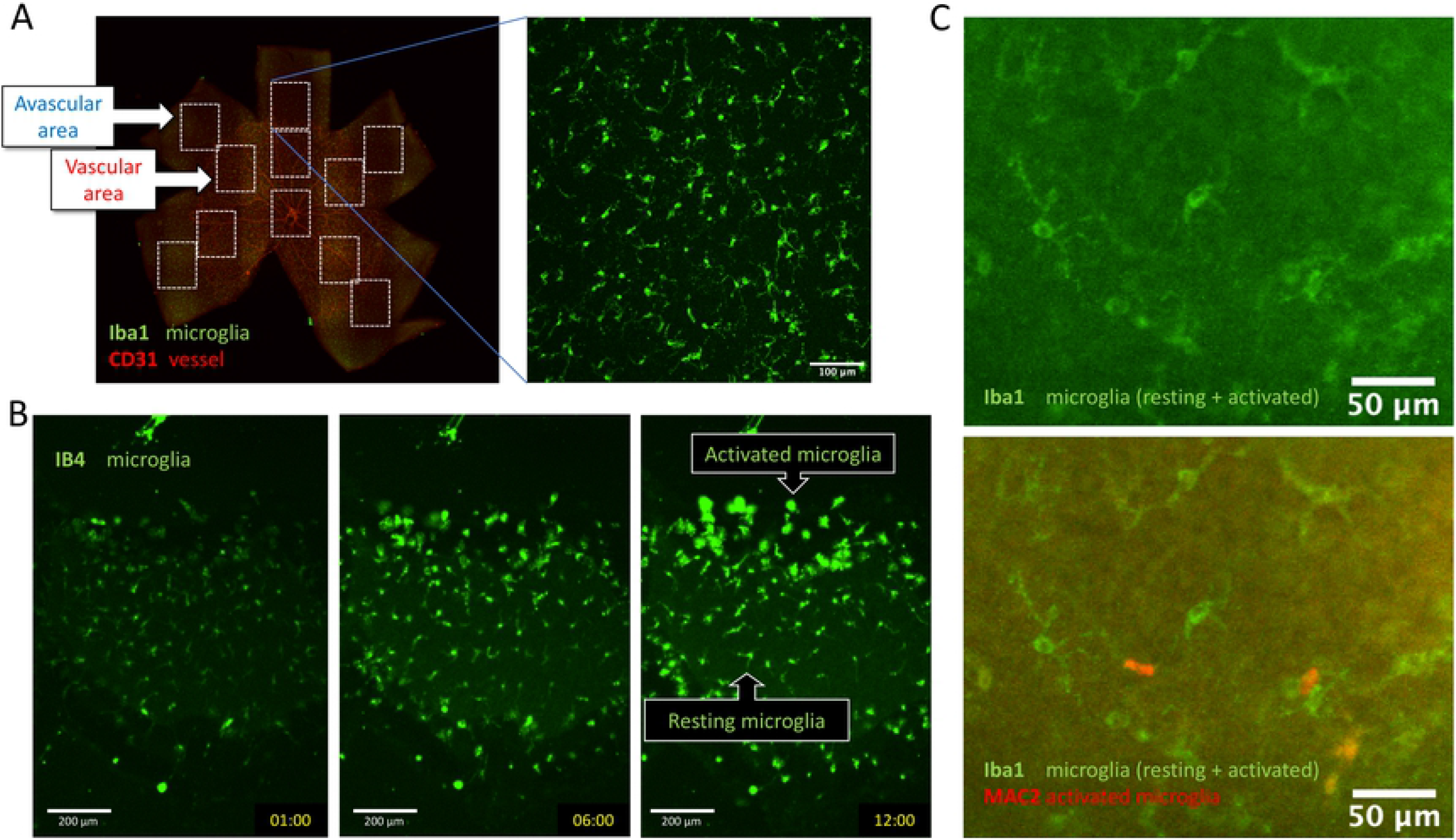
Microglia distribution and morphology. (A) Microglia in the peripheral avascular and the central vascular area of a P5 mouse retina. (B) Observation of activated microglia in the organ culture. A part of the resting microglia was converted to activated microglia and migrated to the injured peripheral edge during a 12-hour time-lapse assay of mouse retina organ culture. (C) Activated microglia stained with MAC2 after the organ culture.

Microglia are derived from primitive hematopoiesis progenitors in the fetal yolk sac during the early embryonic stage [12, 13]. They migrate into all CNS regions, disseminate through the brain parenchyma, and acquire a specific ramified morphological phenotype [14]. Quail microglial precursors enter the retina from the optic nerve head and migrate in a central to peripheral direction. ATP plays a role in the entry and migration of microglial precursors into the developing quail retina. The mechanism for microglial colonization of the CNS appears to be a conserved process across vertebrates [15]. Other adult tissue-resident macrophages are Kupffer cells in the liver, Langerhans cells in the epidermis, and alveolar macrophages in the lung [16].

Advancing microscopy and image processing techniques have made the assessment of microglia morphology more feasible. Nimmerjahn *et al*. reported a highly active motile processes of resting microglial cells using in vivo two-photon imaging [7]. Davis *et al*. reported microglia density, nearest neighbor distance, regularity index, and phenotypes such as soma size and roundness of the mouse retina [17]. However, the mechanism and the molecular pathway of the regular distribution are unclear.

In the present study, we quantified the microglia distribution pattern and confirmed the pattern regularity. Next, we formulated a mathematical model that includes the known regulatory factors of microglial migration. To verify the models experimentally, we quantified the model parameters, reproduced the dynamics with numerical simulations, and tried to specify the molecular pathway responsible for the repulsion between neighboring microglia.

## 3 Materials and methods

### 3.1 Tissue preparation

We used postnatal day five (P5) wild-type mice (Slc: ICR strain) purchased from Japan SLC, Inc., Japan. We sacrificed the P5 mice by decapitation and isolated eyeballs from the orbit for organ culture. After removing eyelids and fat tissues, we made a small hole at the cornea center with a 26G needle. The sclera was gradually removed from the hole using fine forceps, and we isolated the retina in Hanks’ Balanced Salt Solution (HBSS). We removed the lens, vitreous body, and vitelline arteries and prepared the retina fragment with fine forceps. All the animal experiments were undertaken under the permission of the Kyushu University animal experiment committee (A29-036-1).

### 3.2 Immunohistochemistry

We performed immunohistochemistry (IHC) of whole-mount samples or tissue sections as previously described [18]. Eyes were fixed for 20 min in 4% paraformaldehyde (PFA) in phosphate-buffered saline and then dissected. Retinal cups were post-fixed for 30 min and then stained by a standard protocol.

Hamster anti-CD31 (1:1,000; 2H8; Chemicon, Temecula, CA), MAC2 (CEDARLANE), and Iba1 (1:1,000; WAKO, Osaka, Japan) were used as primary monoclonal antibody. For nuclear staining, we treated specimens with DAPI (Molecular Probes) or Hoechst 33342 (Dojindo).

### 3.3 Organ culture

We dissected P5 mouse retinas in HBSS(+). Retinal pieces were cultured on a cell culture insert with a 0.4 µm pore diameter (Millicell PICM0RG50, Merck Millipore Ltd, Cork, Ireland) in DMEM/Ham’s F-12 without Phenol Red (Nacalai Tesque, INC. Kyoto, Japan) supplemented with 10% FBS and 1 % antibiotics (Penicillin-Streptomycin Mixed Solution, Nacalai Tesque, INC. Kyoto, Japan). We added Ib4 lectin (1/1000, Invitrogen I21411) to stain both resting and activated microglia.

Reagents used in the organ culture experiments are: Phosphatidylinositol-specific Phospholipase C from Bacillus cereus (PI-PLC, 0.5 U/ml, Sigma-Aldrich P5542), Recombinant Mouse sFRP-1 Protein (1 µg/ml, R&D Systems, MN9019-SF-025), Plexin-D1 Fc fragment (30 µg/ml), Alexa Fluor 647-ATP (5 µM, Invitrogen A22362), and Clopidogrel (25 µM, TOCRIS 249010).

### 3.4 Extracellular ATP dynamics

We quantified the ATP uptake and diffusion using Alexa Fluor 647-ATP (Invitrogen A22362). We added 5 µM of the fluorescently labeled ATP to the culture medium of the P5 retina organ culture. After 1 hour of incubation, we observed the ATP diffusion of the ATP using fluoresecence recovery after photobleaching (FRAP). The diffusion coefficient is obtained according to [19]. ATP uptake by microglia was assessed by observation of the culture after 12 hours.

We observed the extracellular ATP concentration with ATPOS fluorescent sensor [20, 21]. Dissected P5 retina cups were washed in pH 7.4 HEPES buffer solution (HBS. 25 mM HEPES, 125 mM NaCl, 4 mM KCl, 2 mM CaCl2 and 1 mM MgCl2), and incubated for 5 minutes at room temperature in 100 µl of 0.33 µM neuronal-surface-targeted ATPOS, a molecular complex of ATPOS with BoNT/C-Hc and Alexa Fluor 488 (Alexa488) labeled-streptavidin [20, 21] with 1/100 IB4-DyLight649 (DL-1208-5, Vector Laboratories). Then, the retina cups were washed with HBS + 0.1% BSA three times and transferred on the slide glass of a glass-bottom dish (3911-035-IN, Iwaki Glass). After supplementingg with 500 µl of DMEM/F-12 + 10 % FBS, the optic cups were dissected, and the retina’s anterior half was observed using a Nikon A1 confocal microscope. Extracellular ATP distribution was estimated using the Cy3/Alexa488 signal ratio [20]. We checked the fluorescence response of ATPOS in the organ culture by local application of 2 µl of 10 mM adenylyl-imidodiphosphate (Sigma-Aldrich A2647) (AMP-PNP), an ATP analog that is not degraded by ATPase, as a positive control. The final concentration of AMP-PNP should be 10 µM in the culture medium. The AMP-PNP concentration was calculated using a molar absorption coefficient of 15,000 /M/cm at 260 nm.

### 3.5 Image analysis

#### 3.5.1 Evaluation of microglia morphology

We acquired the confocal images (30 µm z-stack at 3 µm intervals, Nikon A1, 20*×* objective) of the mouse retina. We performed the image processing using ImageJ (Fiji) software [22] (Fig. 2A). First, we created the maximum intensity projections of Z stacks, removed the noise by despeckling and smoothing, and obtained the binary images by thresholding. Since the skeleton length (microglia process) increased as the threshold values decreased, the threshold values were determined manually just before the skeleton length increased explosively (35.2 *±* 0.84, mean *±* SE).

**Fig 2.**
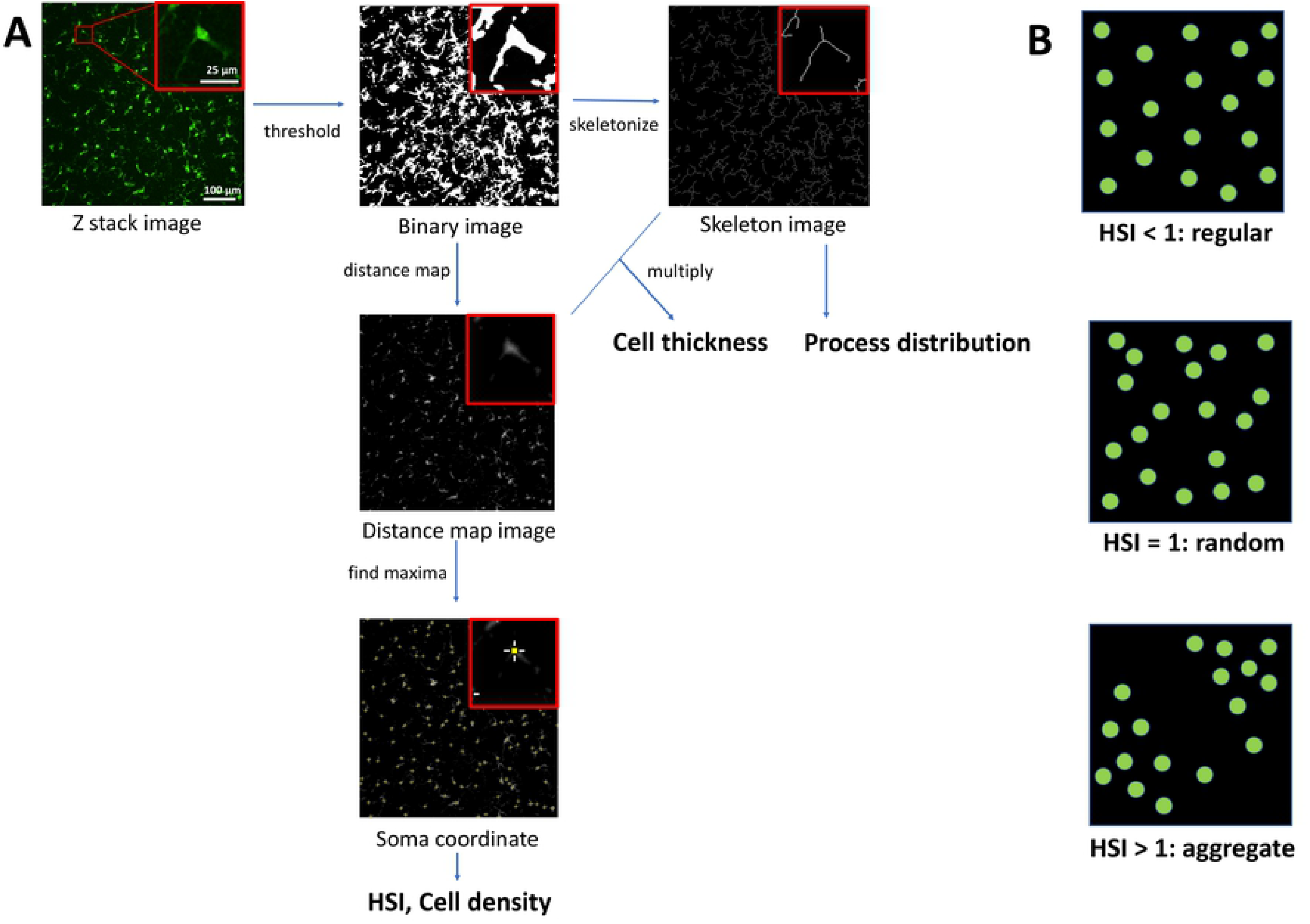
Image Analysis. (A) Image processing for evaluation of microglia morphology and distribution. (B) HSI and regularity of the soma distribution.

Next, we obtained the distance map images and the skeleton images from the binary images. We calculated the process distribution using the skeleton images. We multiplied the distance map image and the skeleton image for the evaluation of microglia thickness. The mean gray value of the multiplication result represents the mean thickness including the soma and the processes.

#### 3.5.2 Evaluation of microglia distribution

The Hopkins-Skellam index (HSI, [23]) was adapted to analyze cell soma distribution (Fig. 2B). HSI is calculated from the following equation:

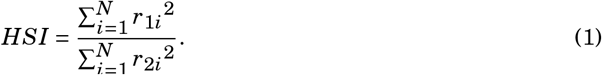

*r*_1_ is a distance from a point chosen at random to its nearest individual, *r*_2_ is a distance from an individual chosen at random to its nearest individual, and *N* is sample size. *HSI <* 1, *HSI =* 1, *HSI >* 1 represents regular, random and aggregate patterns respectively (Fig. 2B).

### 3.6 Numerical simulations

All numerical simulationas are implemented using *Mathematica* (Wolfram Research). The source code of the numerical simulation is provided as an electronic supplementary electronic material.

## 4 Results

### 4.1 Regular distribution of microglia in avascular and vascular area

We quantified the microglia soma distribution and the microglia morphology. We used the P5 mouse retina to observe microglia in both vascular and avascular areas since that microglia are associated with developing vasculature [4]. Microglia are already present in all regions of the mouse retina of mouse embryos aged 11.5 days (E11.5) [24], and angiogenesis starts at birth from the center but does not reach the retinal edges at P5 [25]. We observed 10 avascular (peripheral) and 10 vascular (around the center) areas in two retinae (Fig. 1A).

HSI of both avascular and vascular areas are smaller than 1 (0.56 *±* 0.02 vs. 0.53 *±* 0.02, respectively), indicating that the microglia distribution pattern is regular (Fig. 3 AB). The microglia density is higher in the vascular area (409 *±* 11 vs. 482 *±* 13; P < 0.001). The microglia thickness, including cells and processes, is higher (3.7 *±* 0.5 vs. 2.0 *±* 0.2; P < 0.001) and the average process length is shorter in the avascular area (9.2 *±* 0.08 vs. 9.5 *±* 0.1; P < 0.01), suggesting avascular (peripheral) microglia shows more amoeboid morphology. There was no significant difference in HSI, the number of processes per cell (7.4 *±* 0.8 vs. 7.0 *±* 0.6) and process length per cell (68 *±* 7 µm vs. 66 *±* 5 µm) between avascular and vascular areas (Fig. 3C).

**Fig 3.**
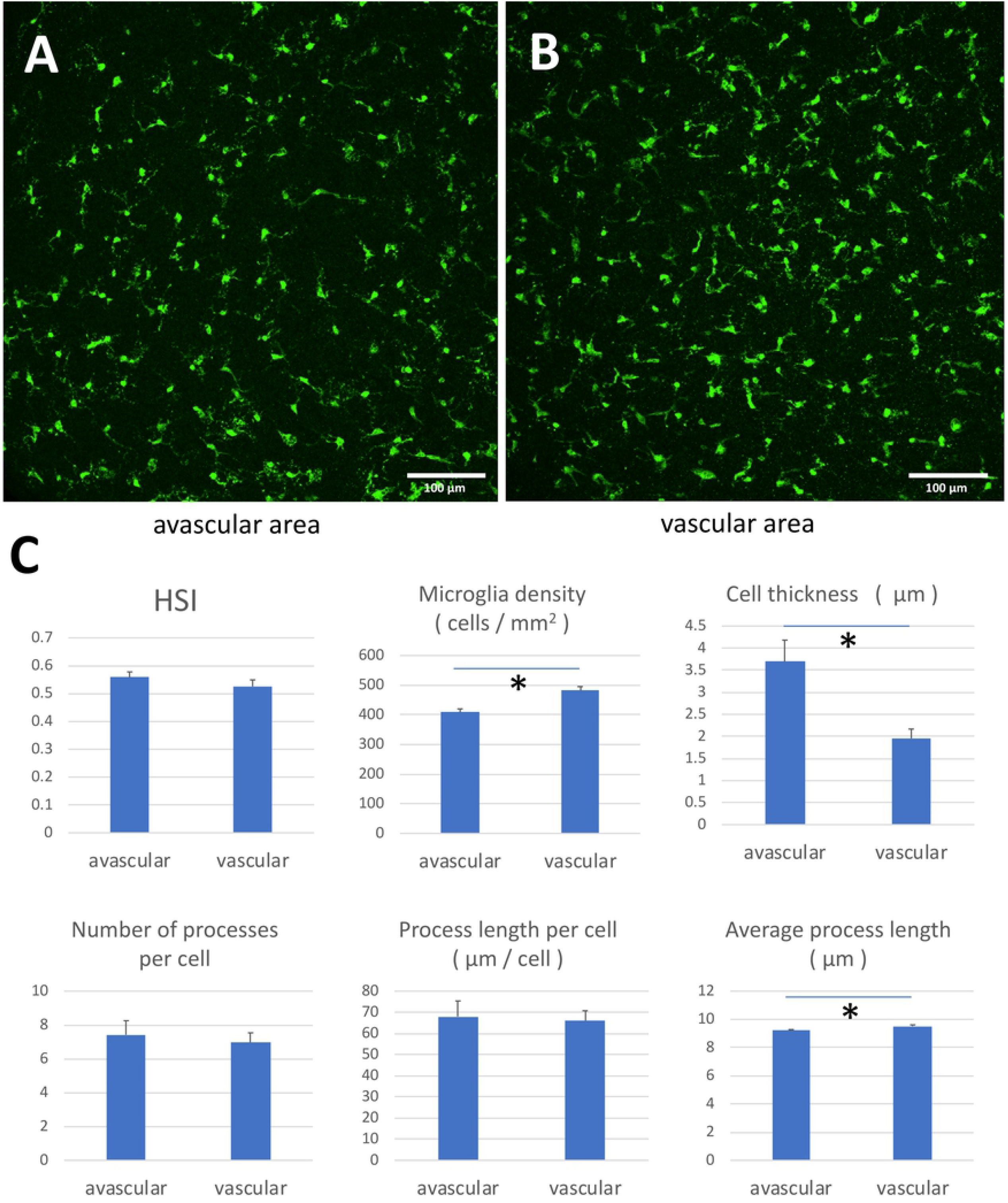
Microglia distribution and morphology. (A) Microglia in the avascular area. (B) Microglia in the vascular area. (C) Microglia density and average process length are higher in the vascular area, while cell thickness is higher in the avascular area. There was no significant difference in HSI, the number of processes per cell, and process length per cell between vascular and avascular areas (Student’s *t*-test). All data are presented as mean *±* SE.

### 4.2 A model of regular microglia distribution

#### 4.2.1 Factors involved in the microglia movement model

Since the critical factor for the observed regular microglia distribution is not clear, we listed up possible factors involved in the microglia distribution.

- **Initial distribution**: In some developmental systems, the cell arrangement pattern is regular from the beginning (like Turing patterns [26] or Delta-Notch system [27]). In the case of microglia, they migrate from the fetal yolk sac to the retina during development [13, 15] and regular spacing is reported only at a late stage of development [1]; thus, we assume that microglia initial distribution is random.
- **Random cell movement**: We observed random cell movement in microglia in organ culture (Fig. 4D). The cells show persistent random walk, which may interfere with regular cell distribution.
- **Chemotaxis toward ATP**: It is well known that ATP is a major chemoattractant of microglia [8–10], which should influence the cell distribution.
- **ATP uptake**: Pinocytosis of microglia has been reported previously [28]. We presumed that when ATP uptake by microglia is combined with ATP chemotaxis, a local decrease of chemoattractant may result in the mutual repulsion of cells [29].
- **Movement by cell-cell contact**: We hypothesized that cell-cell repulsion by direct contact plays a role in establishing the regular distribution of microglia. Although the repulsion by direct contact is not reported in microglia, some cells do repel each other by direct contact (contact inhibition of locomotion, CIL).

**Fig 4.**
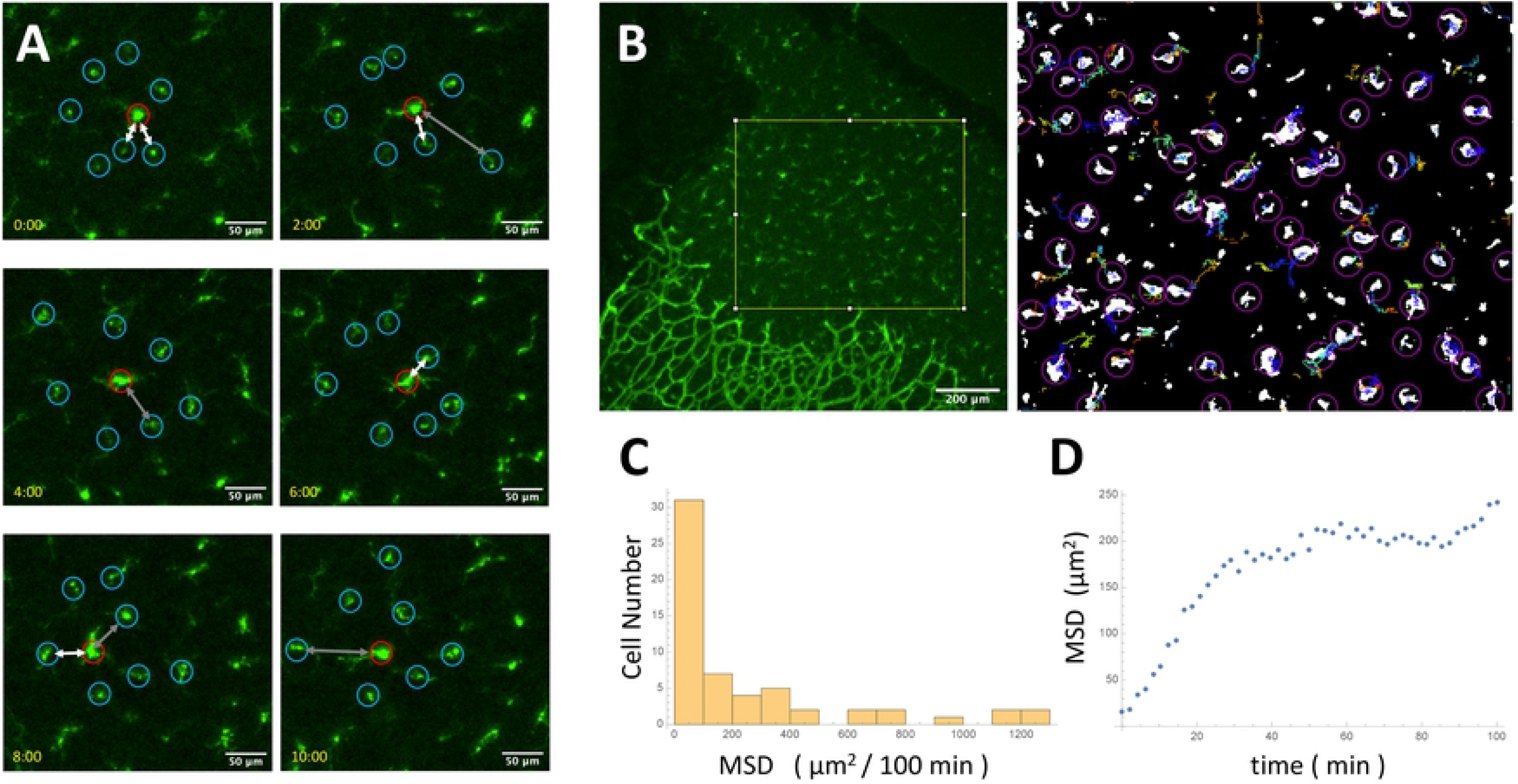
Dynamics of microglial soma in organ culture. (A) Microglia (red circle) moving randomly and repelling surrounding other microglia (blue circles) tracked for 10 hours. White arrows represent the pairs of microglia close to each other, and gray arrows represent the pairs of microglia that move away after approaching. (B) We tracked the microglia migration inside the yellow square in the left panel for 12 hours. The right panel shows that white regions representing each cell’s first position and color lines representing each cell’s 12-hour trajectory. (C) The Histogram of MSD. We chose trajectories we could track for 50 frames (100 min, n = 58). (D) The mean MSD reached a plateau after 40 min.

#### 4.2.2 The model

We propose a model that includes all the factors described in the previous subsection as follows:

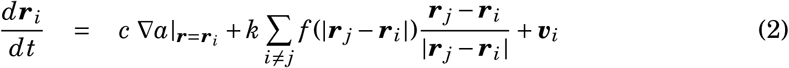

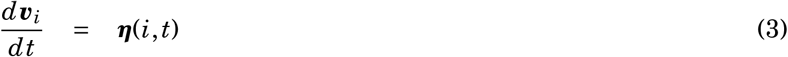

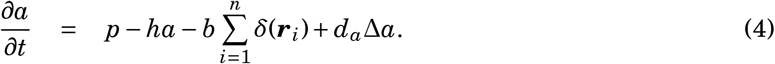

(2) defines the cell movement; ***r***_*i*_ represents the location of the *i*th microglia cell and *a* is the ATP distribution. *c* is ATP chemotactic coefficient and *f* (*r*) is repulsion function (Fig. 7A). When |***r***_*j*_ − ***r***_*i*_| is longer than repulsive radius (*σ*), *f* (|***r***_*j*_ − ***r***_*i*_|) *=* 0. When |***r***_*j*_ − ***r***_*i*_| is shorter than *σ, f* (|***r***_*j*_ − ***r***_*i*_|) *= k*|***r***_*j*_ − ***r***_*i*_|; *k* is microglia repulsive strength coefficient. The function *f* (*r*) mimics that microglia repel each other only when they are close.

(3) implements the persistent random walk. The *i*th cell moves at speed *v*_*i*_ = |***v***_*i*_|. The cell changes the direction of movement at average interval *τ* (persistence), which is implemented by the stochastic term ***η***(*i, t*).

(4) describes the dynamics of ATP concentration; *p* is ATP production rate, *h* is ATP decay coefficient, *b* is ATP uptake rate, and *d*_*a*_ is ATP diffusion coefficient. Microglia cells move according to ATP concentration and internalize ATP (Fig. 6). ATP is generated in the whole domain (*p*), decay (*ha*) and diffuse passively (*d*_*a*_Δ*a*).

### 4.3 Quantification of the model parameters

#### 4.3.1 Dynamics of microglial soma in organ culture

To quantify the dynamics of microglia, we observed microglia in the retinal organ culture. We found that microglia move randomly and appear to repel neighbors when they get closer (Fig. 4A). We tracked the microglia migration (Fig. 4B), and analyzed the tracks that are longer than 50 frames (100 min). We found that most microglia migrate a short distance (MSD *≈* 100 µm^2^ / 100 min, Fig. 4C). A part of the microglia migrated a long distance, which may be activated microglia because of the tissue damage by the dissection of the cultured retinas. We calculated the average of mean square displacement (MSD) with passage time (Fig. 4D); the MSD showed a linear increase initially but reached a plateau after 40 min., indicating that the microglia migration area is limited.

We obtained persistent random walk parameters from the diffusion coefficient (=MSD/time) and average speed (Fig. 4). Diffusion coefficient (*d*_*u*_) and average speed of the microglia (*v*) were *d*_*u*_ *=* 1.9 µm^2^/min and *v =* 0.26 µm/min respectively. Persistence *τ* was calculated by *d*_*u*_ /*v*^2^ *=* 28.1 min.

#### 4.3.2 Dynamics of microglia process formation

An obvious candidate for the cell-cell repulsion mechanism is the microglia processes. We observed the dynamics of microglia process formation using organ culture. We found the processes showed slow growth and rapid collapse (Fig. 5A-C). We measured the average growth velocity of processes *g* (0.36 *±* 0.04 µm / min, mean *±* SE, n = 5) and the collapse probability *p* (0.046 *±* 0.002 / min, mean *±* SE, n = 10). Based on this observation, we made a mathematical model of process distribution that predicts that microglia processes’ length distribution obeys exponential distribution (Supporting Information 6.1).

**Fig 5.**
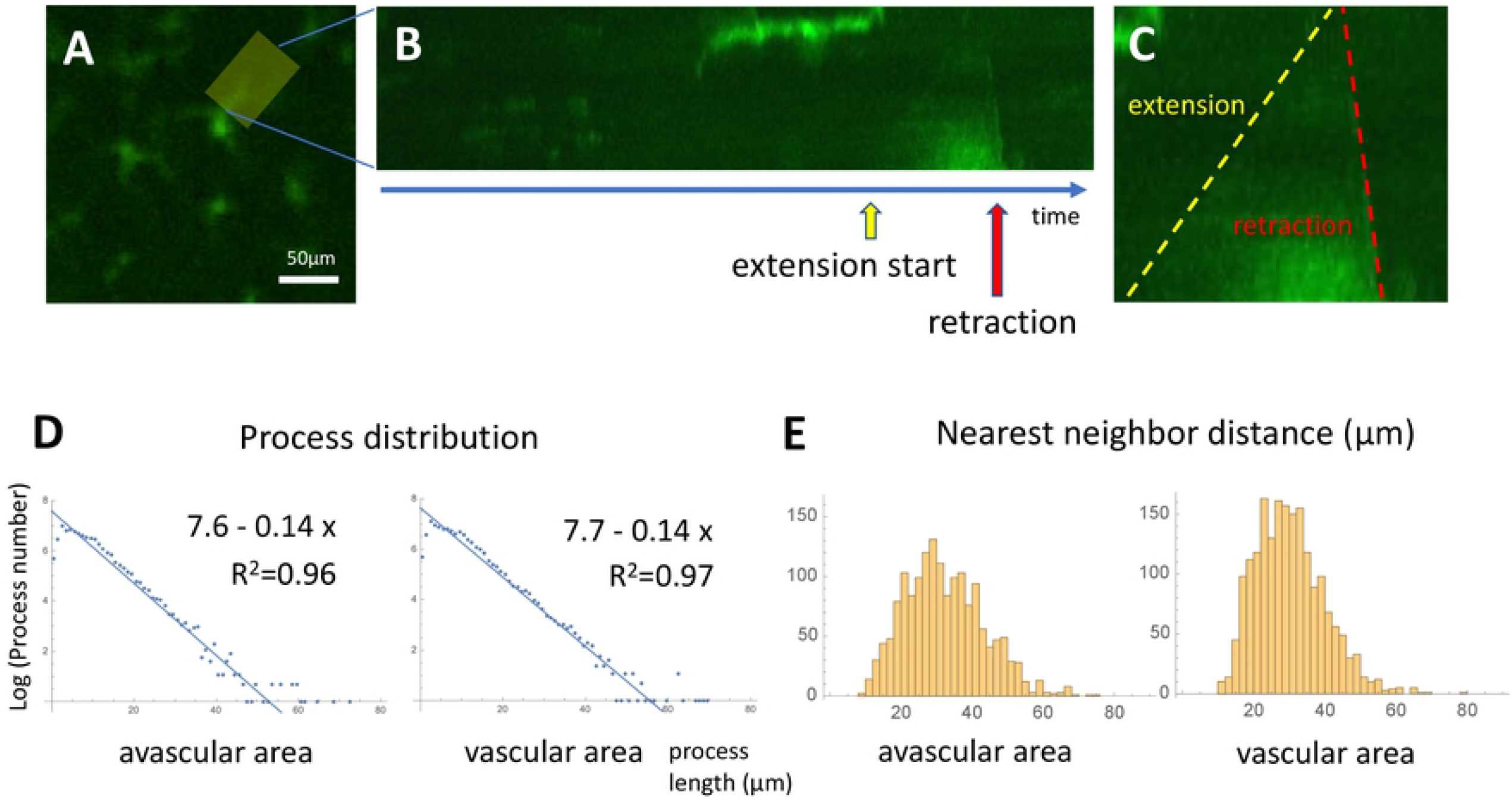
Dynamics of microglia process formation. (A) We obtained the kymograph of microglia process dynamics in the yellow area for 12 hours. Scale bar: 50 µm. (B) Microglia extended the process slowly and retracted quickly. (C) The process extension inclination is the yellow line, and the retraction is the red line. (D) The histogram of process distribution. (E) The histogram of NND.

The experimental results of length distribution of the fixed specimen (Fig. 3) are represented by blue dots (Fig. 5D); the relationship between the log of process number and process length is linear (*y =* 7.6 − 0.14*x, R*^2^ *=* 0.96, avascular area vs. *y =* 7.7 − 0.14*x, R*^2^ *=* 0.97, vascular area), which is consistent with model prediction. The values 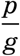 expected from the process dynamics model (0.13) are consistent with the linear fitting slope of the actual process distribution (0.14).

Additionally, we measured the nearest neighbor distance (NND) in both the avascular and the vascular areas. Minimum NND are 9.6 µm vs. 10 µm, and mean NND are 32 *±* 0.3 µm vs. 30 *±* 0.2 µm in avasuclar and vascular areas respectively (mean *±* SE, n = 1619, n = 1910, Fig. 5E). Intuitively, microglia do not get closer beyond the average process length (9.2 *±* 0.08 vs. 9.5 *±* 0.1, mean *±* SE, Fig. 3) because of the direct contact repulsion; therefore, a minimum NND smaller than the average process length does not exist.

#### 4.3.3 ATP uptake, diffusion and extracellular distribution

We also quantified ATP dynamics in relation to microglia in mouse retina organ culture. We previously reported that the lung epithelium exerted a lateral inhibitory effect on the neighboring epithelium via depletion of fibroblast growth factor [29]. We presumed that microglia ingest ATP, decreased the neighborhood’s concentration, and keep regular distribution. We observed microglia distribution and ATP concentration (Alexa Fluor(tm)647 ATP, Invitrogen Corporation, US) using the mouse retina organ culture. We found ATP colocalizes with the cell body of microglia (Fig. 6A), indicating that the uptake of ATP by the microglia does exist. We also measured the change in fluorescence change of microglia in organ culture with 5 µM ATP-Alexa (n = 15) for 12 hours and obtained the uptake rate *b =* 9.5 *×* 10^−4^ µM min^−1^ per microglia cell.

**Fig 6.**
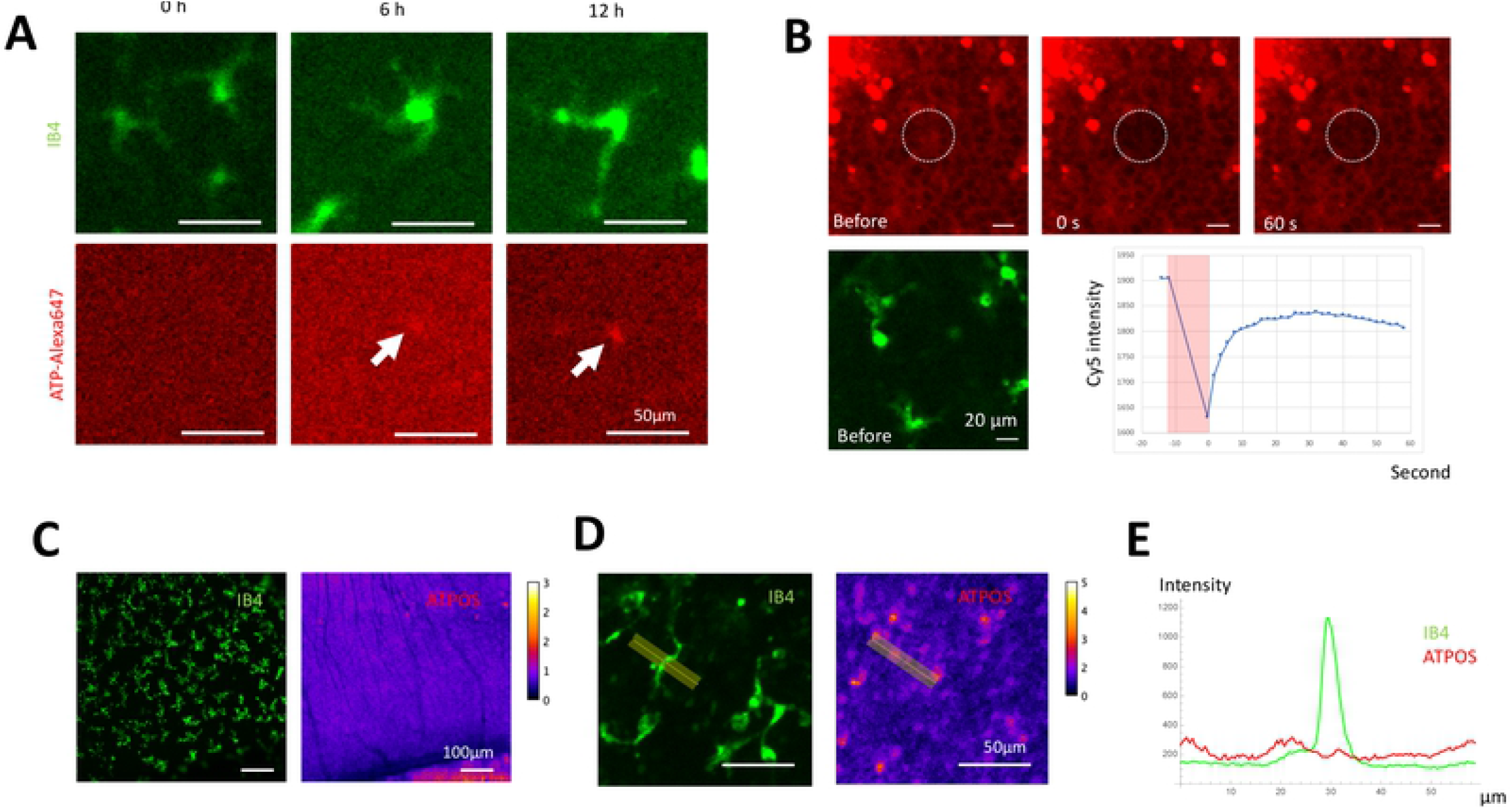
ATP dynamics in mouse retina organ culture. (A) ATP uptake of retinal microglia during 12h. Scale bar: 50 µm (B) Measurement of the ATP diffusion coefficient by FRAP. After the sample was immersed with florescent ATP solution, a small region of the retina was photobleached, and the recovery was observed. (C-E) The extracellular distribution of ATP. (C) Low magnification view. (D) High magnification view. (E) Fluorescence intensity ratio around microglia.

Next, we quantified the diffusion of ATP in the cultured mouse retina using FRAP. Mouse P5 retina was incubated with 50 µM ATP-Alexa647 for 1 hour before the experiment. The dotted circle indicates the bleach spot. From the recovery curve of the fluorescence, we estimated that the diffusion coefficient of ATP is 180 µm^2^ min^−1^. We observed an immobile fraction in the recovery curve (Fig. 6B), indicating that the diffusion of a certain compartment of ATP is very slow [30].

Finally, we observed the extracellular ATP distribution and microglia using ATPOS [20, 21] (Fig. 6C-E). Extracellular ATP was detected using the Cy3-Alexa488 ratio, and the location of microglia was visualized using IB4-DyLight649. We could not detect a decrease in extracellular ATP distribution around the microglia (Fig. 6E).

#### 4.3.4 Model parameters

We evaluated parameter values from the experiments described above to simulate the dynamics of microglia migration and ATP concentration (Table 1). Some parameters are estimated from previous work, which is described in Supporting information (section 6.2).

**Table 1.** Simulation parameter values in the model

| Parameter | Description | Value | Source |
| --- | --- | --- | --- |
| $c$ | ATP chemotactic coefficient | $18 \mu\text{m}^2 \mu\text{M}^{-1} \text{min}^{-1}$ | [8] |
| $\tau$ | Persistence | 11.6 min | Fig. 4 |
| $v$ | Random cell migration speed | $2.5 \mu\text{m} / \text{min}$ | Fig. 4 |
| $p$ | Extracellular ATP production rate | $1.7 * 10^{-3} \mu\text{M} \text{min}^{-1}$ | Calculated from (4) |
| $h$ | ATP decay coefficient | $0.2 \text{min}^{-1}$ | [31] |
| $b$ | ATP uptake rate | $9.5 * 10^{-4} \mu\text{M} \text{min}^{-1} \text{microglia}^{-1}$ | Fig. 6 |
| $d_a$ | ATP diffusion coefficient | $180 \mu\text{m}^2 \text{min}^{-1}$ | Fig. 6 |
| $d_u$ | Microglia diffusion coefficient | $0.4 \mu\text{m}^2 \text{min}^{-1}$ | [32] |
| $\sigma$ | Microglia repulsive radius | 45 $\mu\text{m}$ | Fig. 5 |
| $k$ | Repulsive strength coefficient of discrete model | $0.1 \text{min}^{-1}$ | Determined by simulation |
| $a_w$ | ATP concentration at wound site | 31 $\mu\text{M}$ | Determined by simulation |

### 4.4 The numerical simulations of the model

We numerically confirmed whether the parameter set we obtained is sufficient to generate regular spacing.

Among the model parameters, the repulsion strength coefficient *k* is the only parameter that the experimental observation cannot determine. We numerically obtained the effect of the different *k* to the regular distribution of the microglia cells. *k* works very efficiently in terms of the regular pattern formation, and small parameters like *k =* 0.1 can generate a regular pattern (Fig. 7A). *k* represents the inverse of the time required for the repulsion to take effect. At first, we assumed that the depletion of ATP could also play a role, but *k =* 0 simulation shows no regular distribution, indicating the effect of direct cell-cell repulsion may be the primary factor.

**Fig 7.**
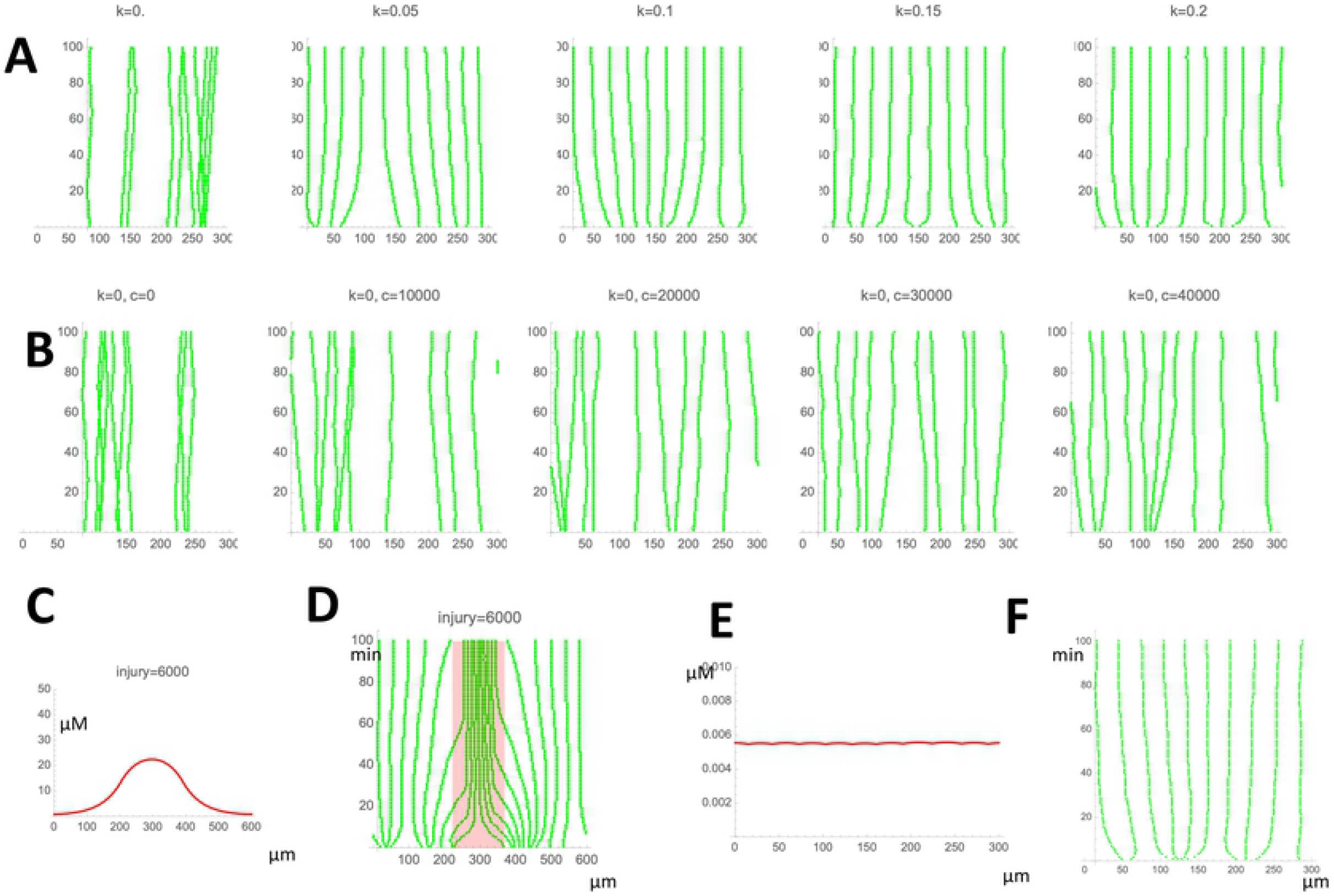
The model simulations. (A) Determination of *k*. (B) Effect of chemotactic coefficient *c* when *k =* 0. (C) Distribution of ATP at the wound site in the model. (D) Accumulation of microglia at the wound site in the model. (E) Distribution of ATP in the normal tissue in the model. (F) Microglia distribution of the quantitative model.

One concern about the parameter estimation and ATP effect is that the chemotactic coefficient *c* may be underestimated because it is based on an in vitro experiment [8]. To rule out this possibility, we first observed what *c* value can reproduce regular spacing without cell-cell repulsion (*k =* 0). We needed a very large chemotactic factor for the regular spacing (*c =* 20000) from the model simulation, which is 1000 times higher than the observed value. In addition, the original model parameter (*c =* 18) reproduced the accumulation of the microglia cells at the wounded site (Fig. 1B). Since the production rate of the ATP at the wound site is unknown, we set the concentration of the ATP at the wound site as the 1/1000 of the intracellular concentration (Fig. 7c, [33]). We observed the accumulation of the microglia cells at the wound site (Fig. 7D, red), indicating that the chemotactic coefficient *c* is appropriate. The final ATP distribution and microglia cell distribution are shown in Fig. 7 EF. We observed the decrease of ATP concentration at the microglia location, but the concentration decrease is very small (Fig. 7E), consistent with ATPOS observation (Fig. 6C-E).

### 4.5 Screening molecular pathways for cell-cell interaction

Finally, we tried to specify the molecular pathway responsible for the direct contact repulsion or ATP. Since the mechanism of repulsion by cell-cell contact has not been described in microglia, we screened three pathways responsible for CIL: the ephrin-Eph pathway, the Wnt-planar cell polarity (PCP) pathway, and the Semaphorin pathway.

We confirmed Eph-ephrin, Wnt-PCP, and semaphorin pathway gene are expressed in mouse microglia using RefEx (Reference Expression dataset) [34]. We used phosphatidylinositol-specific phospholipase C (PI-PLC) which inhibits the EphA-ephrinA system, Secreted Frizzled Related Protein 1 (sFRP1) to block the Wnt-PCP pathway [35], and Plexin-D1 Fc fragment to inhibit the semaphorin pathway.

We observed microglia distributions at 400 µm *×* 400 µm in the avascular area of organ culture for 12 hours (Fig. 8A), treated with PI-PLC, sFRP1, and Plexin-D1 Fc fragment assumed to inhibit microglia cell-cell repulsion. THe HSI of the treated groups tends to be higher than the control (Fig. 8B), indicating that inhibitors of cell-cell repulsion interfere with regular microglia distribution. Microglia densities of the treated groups are lower than the control (Fig. 8C). We assume that microglia migrate at ease because of a lack of surrounding repulsion; thus, individual microglia migrate freely and into the peripheral damaged area. Therefore the cell density decreases within the observed field.

**Fig 8.**
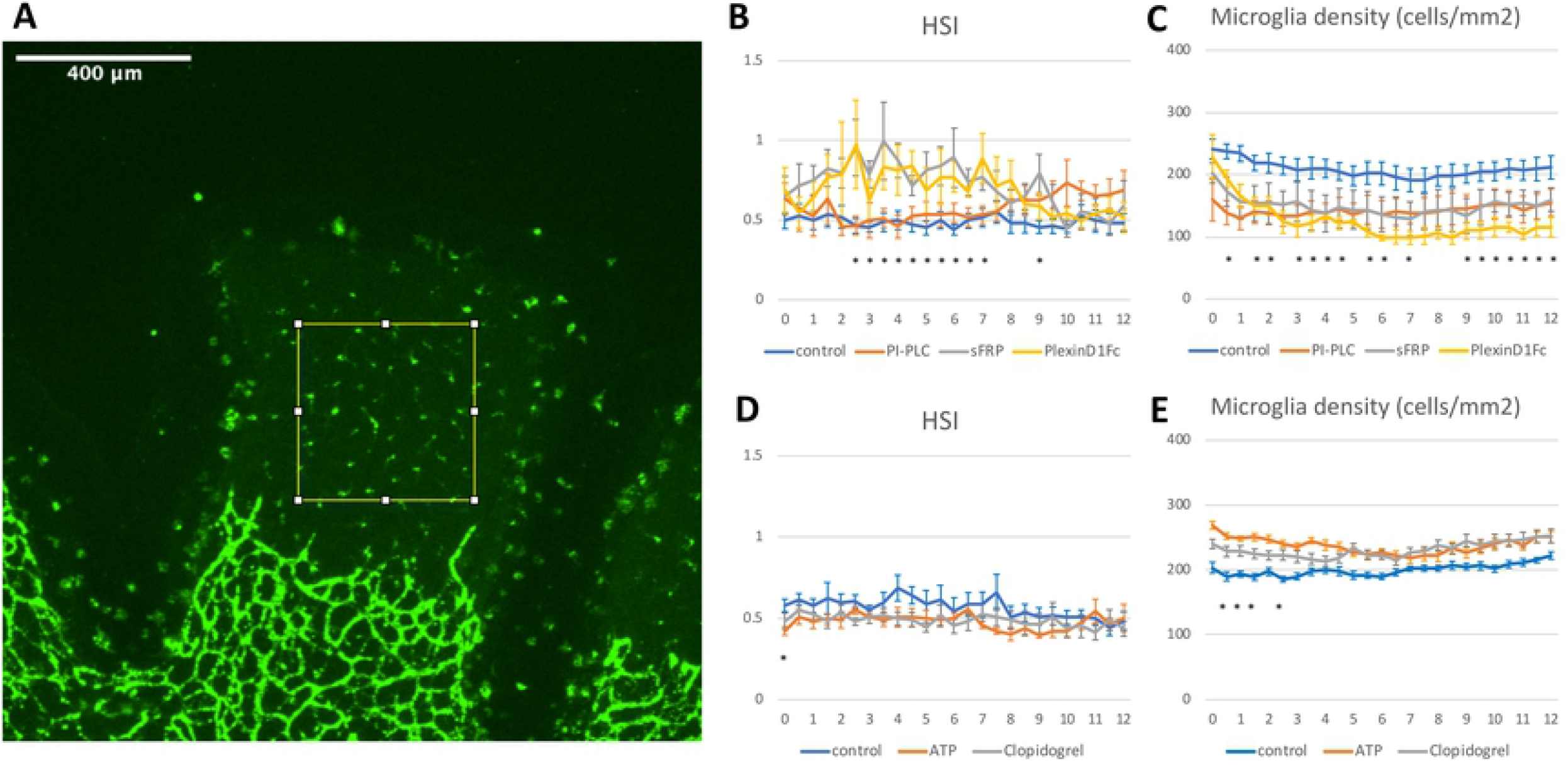
HSI and cell densities of microglia in organ culture. (A) 400 µm *×* 400 µm areas were measured. (B) HSI of the repulsion assay (control: *n =* 7, PI-PLC: *n =* 5, sFRP: *n =* 5, PlexinD1Fc: *n =* 4). (C) Cell densities of the repulsion assay. (D) HSI of the ATP assay (control: *n =* 7, ATP: *n =* 8, Clopidogrel: *n =* 7). (E) Cell densities of the ATP assay. All data are represented as mean *±* SE. *: one-way ANOVA, *P <* 0.05.

Besides, we observed the microglia distribution treated with a saturating concentration of ATP-Alexa and clopidogrel (ATP P2Y receptor antagonist). The HSI of treated groups tends to be similar to that of the control group (Fig. 8D). Microglia densities of treated groups are higher than control (Fig. 8E). We assume that saturating amount of ATP or ATP inhibitor mask ATP gradient from the peripheral damaged area to the center of observed tissue; thus, treated microglia stay in the same position.

## 5 Discussion

### 5.1 The biological meaning of regular distribution

The physiological meaning of the regular distribution is not clear. The structure of resting microglia serves an immune surveillance function [7]. The regular distribution is assumed to be essential for the immune system and enables microglia to reach the injured site.

The regular distribution of microglia may play a role in the pattern formation of blood vessels. It is known that microglia influences angiogenesis [36, 37]. Reduced number of microglia results in reduced vessel branching and density, while patterning was restored by intravitreal injection of exogeneous microglia [38–40].

### 5.2 Models of microglia cell distribution

There are several theoretical studies concerning the distribution of microglia. Numahara *et al*. reported that the distribution of epidermal Langerhans cells is completely regular; the pattern of Voronoi divisions fits the territories hypothesized for the immuno-surveillance. They estimated a repulsive interaction between the Langerhans cells using the mathematical model [41]. However, the mechanism of the regular distribution of Langerhans cells is not specified.

Another model concentrates on aggregation formation of microglia around amyloid-beta plaques, not regular spacing [42]. In this work, aggregation of microglia was reproduced using the Kelle-Segel type chemotaxis model [43].

### 5.3 Microglia process dynamics

Several features are known about the dynamics of process formation. Resting microglia extends the process to the injured site [7] which is mediated by Gi-coupled P2Y12 ATP receptor and downstream Rac GTPase-driven actin polymerization [8]. Process retraction during the transition from resting to active microglia is mediated by the adenosine A_2*A*_ receptor [44]. The speed of process extension and retraction were similar in the brain [7], which is different from our observation in the retina (Fig. 5 C). Resting microglia process movement is interspersed with a brief static period. The physiological meaning of this fixed period is not clear [7].

We investigated the dynamics of the microglia process formation to obtain the characteristic length of repulsion. The mathematical model contains an advection term, representing constant growth of the processes, and degradation term, which represents the probabilistic process collapse. This model fits the experimentally observed process length distribution. This model is similar to our previous model [45] and essentially equivalent to the McKendrick-von Forester equation with a uniform death rate [46]. We found that microglia do not get closer beyond the average process length. We presume that microglia maintain distances between other microglia while extending and retracting the processes.

### 5.4 Contact Inhibition of Locomotion pathway

To our knowledge, there is no previous report describing the mechanism of microglia repulsion via contact by processes. Therefore we examined a general cell repulsion mechanism known as the contact inhibition of locomotion, initially described in the behavior of fibroblast cells [47, 48].

Three molecular pathways of CIL are known - the Eph-ephrin pathway [49], the Wnt-PCP pathway [49, 50] and the Semaphorin pathway [51]. Eph and ephrin are membrane-bound receptors and ligands, respectively, and when Eph-expressing cells contact an ephrin-expressing cell, the cells repel each other. Ephrin-Eph signaling plays an essential role in cell segregation during development [52, 53]. Noncanonical Wnts (e.g., Wnt4, Wnt5A, and Wnt11) activate PCP [54] that is related to CIL. sFRP is one of the extracellular inhibitors of this pathway [35]. Semaphorins are secreted or transmembrane proteins that mediate repulsive axon guidance, immune cell regulation, and vascular growth and remodeling [55]. Semaphorins signal through plexins that recruit and regulate kinases and Rho-family GTPases to control cell motility [56].

### 5.5 Cell-cell repulsion by direct contact in other systems

Although the mechanism of cell-cell repulsion in microglia has not been reported, previous works described the mechanism in other cell types. Mouse oligodendrocyte precursor cells (OPC) survey their local environment with motile filopodia and maintain unique territories through self-avoidance [57]. In *Drosophila* da sensory neurons, the mutual repulsion of dendrites is regulated by gene encoded in Dscam (Down syndrome cell adhesion molecule) locus [58]. PVD nociceptive neuron in C. elegans extends dendrites that do not cross each other. The axon guidance protein UNC-6/Netrin is required for this self-avoidance [59]. Two mouse retinal interneuron subtypes (starburst amacrine cells and horizontal cells) show regular distribution, and MEGF10 and MEGF11 are involved in this regular distribution [60].

## 6 Supporting information

### 6.1 Model of microglia process length distribution

Based on the experimentally observed motion of the process, we modeled the dynamics of process length distribution *u*(*x, t*) using advection and degradation terms as follows:

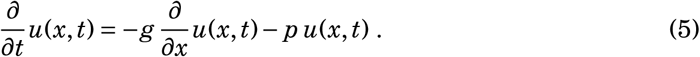

*g* is the velocity of process extension and *p* is the collapse probability of process. By the model (5), the steady-state of length distribution *u*(*x, ∞*) can be described as follows:

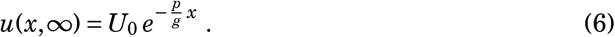

*U*_0_ represents *u*(0, *∞*), the steady-state length distribution of the microglia process. *U*_0_ is exponential distribution, indicating there is a linear relationship between the log of process number and process length.

The sum of the process length is described by the definite integral as follows:

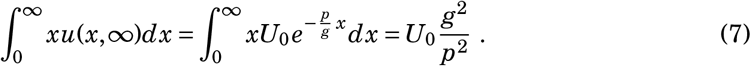

*U*_0_ in avascular are (7.6) and vascular areas (7.7) are obtained by linear fitting equation. 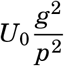 are 1.0 *×* 10^5^ µm vs. 1.1 *×* 10^5^ µm, and the sum of the process length from experimental results are 1.1 *×* 10^5^ µm vs. 1.3 *×* 10^5^ µm, respectively. The model captures the characteristics of the actual process distribution.

### 6.2 Obtaining simulation parameters from previous literatures

- *c*: We calculated the ATP chemotaxis coefficient using the published data of ATP chemotaxis of microglia in the Dunn chamber [8].
- *h*: We determined the ATP decay coefficient by fitting the data that ATP decay of homogenized retina [31]. The fitting equation as follows:

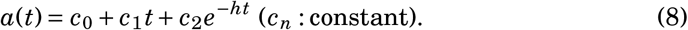
- *p*: The basal extracellular ATP production rate [61] is calculated by steady state concentration of ATP (*a*_0_) and ATP decay rate *h*. We used extracellular ATP of the bovine retina (6.2 *±* 0.7 nM [62]) as *a*_0_. We obtain *p* by steady state equilibrium 0 *= p* − *ha*_0_. Uptake by microglia is small and neglected in this calculation.
- *σ*: Davis *et al*. reported the exclusion radii (58.6 *±* 1.4 µm) and the average NND (40 *±* 1.0 µm) in adult mouse retina [17]. We measured the average NND (31 *±* 0.12 µm, n = 7053, Fig. 3) in P5 mouse retina; thus, we assumed microglia repulsion radius is 45 µm, considering the ratio of their report [17].
- *aw*: Intracellular ATP is millimolar order [33]. We set production of ATP at the wound site so that *aw* becomes 1/1000 of intracellular ATP.

## 7 Acknowledgments

This work is financially supported by JSPS KAKENHI (Grant Nos. 20H03427 to Kenzo Hirose, and 20H02875 to Daisuke Asanuma). The authors would like to thank Shuji Ishihara (University of Tokyo) for discussions and comments, and Enago (www.enago.jp) for the English language review.

## Notes

### Competing Interest Statement

The authors have declared no competing interest.

